# Universal surface-to-volume scaling and aspect ratio homeostasis in rod-shaped bacteria

**DOI:** 10.1101/583989

**Authors:** Nikola Ojkic, Diana Serbanescu, Shiladitya Banerjee

## Abstract

Rod-shaped bacterial cells can readily adapt their lengths and widths in response to environmental changes. While many recent studies have focused on the mechanisms underlying bacterial cell size control, it remains largely unknown how the coupling between cell length and width results in robust control of rod-like bacterial shapes. In this study we uncover a universal surface-to-volume scaling relation in *Escherichia coli* and other rod-shaped bacteria, resulting from the preservation of cell aspect ratio. To explain the mechanistic origin of aspect-ratio control, we propose a quantitative model for the coupling between bacterial cell elongation and the accumulation of an essential division protein, FtsZ. This model reveals a mechanism for why bacterial aspect ratio is independent of cell size and growth conditions, and predicts cell morphological changes in response to nutrient perturbations, antibiotics, MreB or FtsZ depletion, in quantitative agreement with experimental data.

## Main Text

Cell morphology is an important adaptive trait that is crucial for bacterial growth, motility, nutrient uptake, and proliferation [1]. When rod-shaped bacteria grow in media with different nutrient availability, both cell length and width increase exponentially with growth rate [2, 3]. At the single-cell level, control of cell volume in many rod-shaped cells is achieved via an *adder* mechanism, whereby cells elongate by a fixed length per division cycle [4–8]. A recent study has linked the determination of cell size to a condition-dependent regulation of cell surface-to-volume ratio [9]. However, it remains largely unknown how cell length and width are coupled to regulate rod-like bacterial shapes in diverse growth conditions [10, 11].

Here we investigated the control of cell surface area (*S*) relative to cell volume (*V*) for *E. coli* cells grown under different nutrient conditions, challenged with antibiotics, protein overexpression or depletion, and single gene deletions [3, 9, 12–14]. Collected surface and volume data span two orders of magnitude and exhibit a single power law in this regime: *S* = *µV*^2/3^ (Fig. 1A). Specifically, during steady-state growth [3] *µ* = 6.24*±*0.04, suggesting an elegant geometric relation: *S* ≈ 2*πV*^2/3^. This universal surface-to-volume scaling with a constant prefactor, *µ*, is a consequence of tight control of cell aspect ratio *η* (length/width), whose mechanistic origin has been puzzling for almost half a century [15, 16]. Specifically, for a sphero-cylindrical shape, *S* = *µV*^2/3^ implies 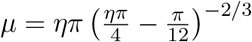. A constant *µ* thus defines a constant aspect ratio *η* = 4.14 *±* 0.17 (Fig. 1B-inset), with a coefficient of variation *∼* 14% (Fig. 1B).

**FIG. 1.**
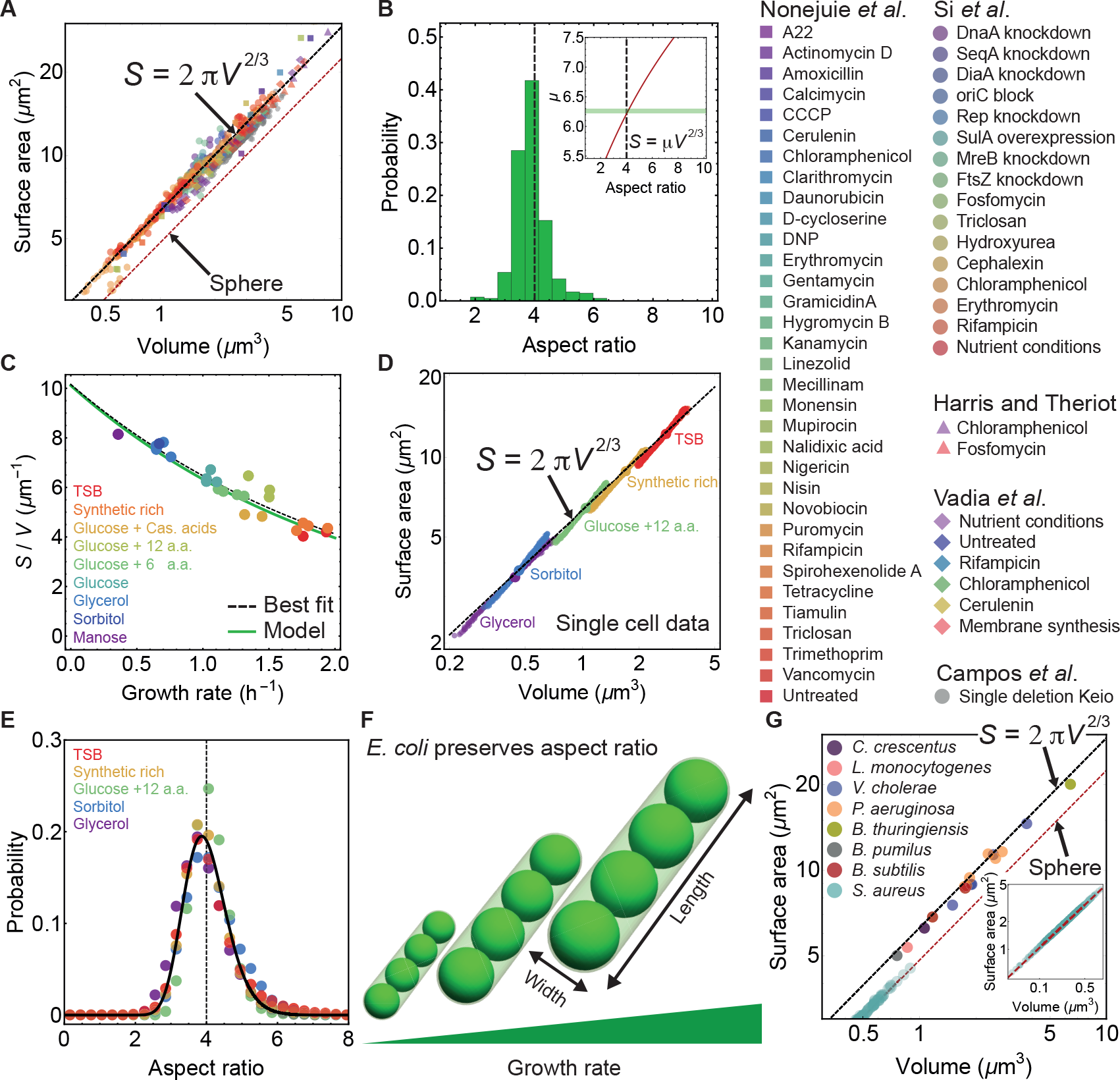
Surface-to-volume scaling in *E. coli* and other bacterial species. (A) *E. coli* cells subjected to different antibiotics, nutrient conditions, protein overexpression/depletion, and single gene deletions [3, 9, 12–14], follow the universal relation between population-averaged surface area (*S*) and volume (*V*): *S* = *µV* ^2/3^ (4981 data points; Supplementary Table 1). Best fit shown in dashed black line for steady-state data from [3] gives *µ* = 6.24 *±* 0.04, and a power law exponent 0.671 *±* 0.006. For single deletion Keio set [14], the best fit curve is *S* = 5.79 *V* ^2/3^. (B) Cell-to-cell variability in aspect ratio [3]. (Inset) Analytical relation between *µ* and aspect ratio *η* for a sphero-cylinder (red line). Best fit from (A) shown with horizontal green band gives aspect ratio 4.14 *±* 0.17. (C) *S/V* vs growth rate. Model line assumes the relation *S* = 2*πV* ^2/3^ and the nutrient growth law (Eq. 1). Data from [3]. (D) *S* vs *V* for newborn *E. coli* cells grown in mother machine [6]. For sample size refer to Supplementary Table 1. Single cell data (small circles) binned in volume follow population averages (large circles). (E) Probability distribution of newborn cell aspect ratio is independent of growth rate, fitted by a log-normal distribution (solid line). (F) Schematic of small, medium, and large *E. coli* cells with a conserved aspect ratio of 4. (G) *S* vs *V* for different bacterial species [9, 17–23]. (Inset) Coccoid *S. aureus* exposed to different antibiotics. Best fit: *S* = 4.92 *V* ^2/3^.

This universal surface-to-volume relationship for steady-state growth results in a simple expression for cell surface-to-volume ratio: *S/V ≈* 2*πV* ^−1/3^. Using the phenomenological nutrient growth law *V* = *V*_0_*e*^*ακ*^[2], where *κ* is the population growth rate, we predict a negative correlation between *S/V* and *κ*:

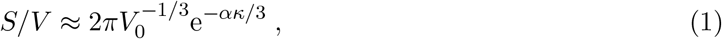

with *V*_0_ the cell volume at *κ* = 0, and *α* is the relative rate of increase in *V* with *κ* (Fig. 1C). Eq. (1) underlies an adaptive feedback response of the cell — at low nutrient conditions, cells increase their surface-to-volume ratio to promote nutrient influx. Prediction from Eq. (1) is in excellent agreement with the best fit to the experimental data. Furthermore, a constant aspect ratio of *≈* 4 implies 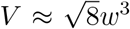 and *S ≈* 4*πw*^2^, where *w* is the cell width, suggesting stronger geometric constraints than recently proposed [11, 24]. Thus, knowing cell volume as a function of cell cycle parameters [3] we can directly predict cell width and length in agreement with experimental data (Figure 1—figure supplement 1A-B).

To investigate how surface-to-volume ratio is regulated at the single cell level we analysed the morphologies of *E. coli* cells grown in the mother machine [6] (Fig. 1D). For five different growth media, mean volume and surface area of newborn cells also follow the relationship *S* = 2*πV* ^2/3^, suggesting that a fixed aspect ratio is maintained on average. Deviation from the mean trend is a consequence of intergenerational or cell-to-cell variabilities in length and width (Figure 1—figure supplement 1C-D). Importantly, the probability distribution of aspect ratio is independent of the growth media (Fig. 1E), implying that cellular aspect ratio is independent of cell size as well as growth rate (Fig. 1F). We further analysed cell shape data for seven additional rod-shaped and one coccoid bacteria (Fig. 1G). Surprisingly, all rod-like cells follow the same universal surface-to-volume scaling, while the coccoid *S. aureus* maintains a much lower aspect ratio *η* = 1.38 *±* 0.18 [21]. This suggests that *aspect ratio homeostasis* likely emerges from a mechanism that is common to diverse bacterial species.

To elucidate the origin of aspect ratio homeostasis we developed a quantitative model for cell shape dynamics that accounts for the coupling between cell elongation and the accumulation of cell division proteins FtsZ (Fig. 2A). *E. coli* and other rod-like bacteria maintain a constant width during their cell cycle while elongating exponentially in length *L* [6]: d*L/*d*t* = *kL*, with *k* the elongation rate. Cell division is triggered when a constant length is added per division cycle - a mechanism that is captured by a model for threshold accumulation of division initiator proteins, produced at a rate proportional to cell size [18, 25, 26]. While many molecular candidates have been suggested as initiators of division [27], a recent study [28] has identified FtsZ as the key initiator protein that assembles a ring-like structure in the mid-cell region to trigger septation.

**FIG. 2.**
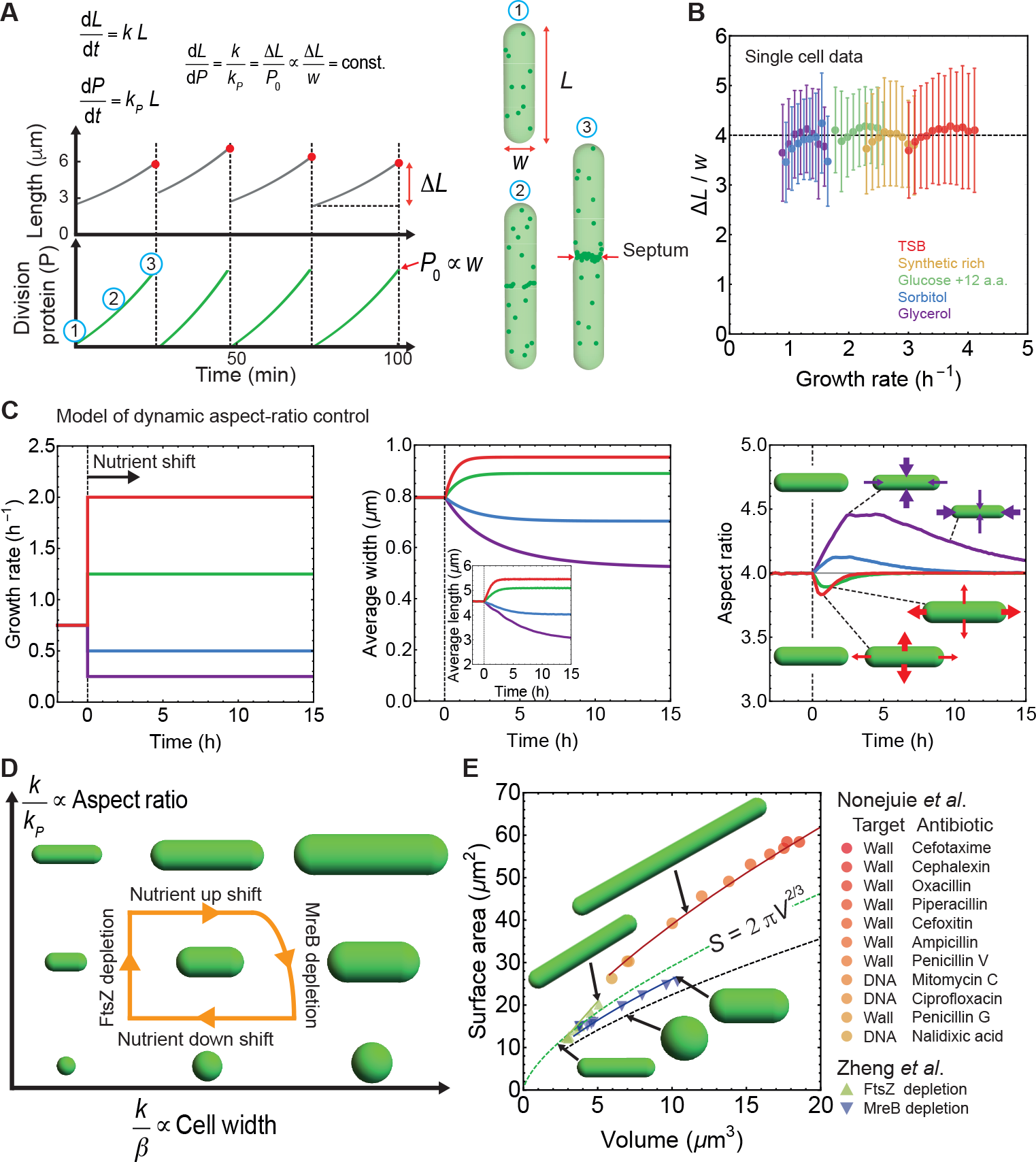
Model for aspect ratio homeostasis in rod-like bacteria. (A) Model schematic. Cell length *L* increases exponentially during the division cycle at a rate *k*. Division proteins (*P*) are produced at a rate *k*_*P*_, and assembles a ring in the mid-cell region. Cells divide when *P* = *P*_0_ *∝ w*. *P* vs time and *L* vs time are reproduced from model simulations. (B) Ratio of the added length (∆*L*) and cell width (*w*) is constant and independent of growth rate [6]. Error bars: *±*1 standard deviation. (C) Simulations of cell-shape dynamics under different growth conditions. (Left) At *t* = 0 h cells are exposed to nutrient upshift or downshift. (Middle) Population-averaged cell length and width vs time. (Right) Population-averaged aspect ratio of newborn cells vs time. (D) Model parameters that control changes in cell aspect ratio (*k/k*_*P*_) or width (*k/β*). (E) Surface area vs volume for cells under antibiotic treatement [12], FtsZ knockdown and MreB depletion [29]. Solid lines are best fit obtained using our model (see Appendix). Cells with depleted FtsZ have elongated phenotypes, while depleted MreB have smaller aspect ratio and larger width. Cell wall or DNA targeting antibiotics induce filamentation. Dashed green line: *S* = 2*πV* ^2/3^, dashed black line: spheres.

Dynamics of division protein accumulation can be described using a two-component model (Appendix). First, a cytoplasmic component with abundance *P*_c_ grows in proportion to cell size (*∝ L*), as ribosome content increases with cell size [30]. Second, a ring-bound component, *P*_r_, is assembled from the cytoplasmic pool at a constant rate. At the start of the division cycle, *P*_c_ = *P*^∗^ (a constant), *P*_*r*_ = 0, and the cell divides when *P*_r_ reaches a threshold amount, *P*_0_, required for the completion of ring assembly. A key ingredient of our model is that *P*_0_ scales linearly with the cell circumference *πw*, preserving the density of FtsZ in the ring. This is consistent with experimental findings that the total FtsZ scales with the cell width [31]. Accumulation of division proteins, *P* = *P*_c_ + *P*_r_ *− P*^∗^, follows the equation: d*P/*d*t* = *k*_*P*_*L*, where *k*_*P*_ is the production rate of division proteins, with *P* = 0 at the start of the division cycle and *P* = *P*_0_ at cell division (Fig. 2A). As a result, during one division cycle cells grow by adding a length ∆*L* = *P*_0_*k/k*_*P*_, which equals the homeostatic length of newborn cells. Furthermore, recent experiments suggest that the amount of FtsZ synthesised per unit cell length, d*P/*d*L*, is constant [28]. This implies,

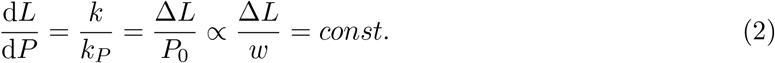

Aspect ratio homeostasis is thus achieved via a balance between the rates of cell elongation and division protein production, consistent with observations that FtsZ overexpression leads to minicells and FtsZ depletion induces elongated phenotypes [29, 32]. Indeed single cell *E. coli* data [6] show that ∆*L/w* is constant on average and independent of growth conditions (Fig. 2B, Figure 2—figure supplement 1A). Thus cell width is a good predictor for added cell length.

To predict cell-shape dynamics under perturbations to growth conditions we simulated our single-cell model (Fig. 2A, Appendix) with an additional equation for cell width that we derived using the model [9]: d*S/*d*t* = *βV*, where *β* is the rate of surface area synthesis relative to volume (Figure 2—figure supplement 1B). For a sphero-cylinder shaped bacterium, we have

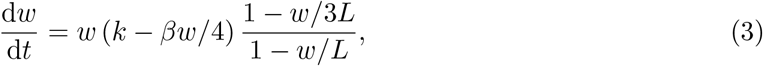

such that *w* = 4*k/β* at steady-state. When simulated cells are exposed to new nutrient conditions (Figure 2—figure supplement 1C-E), changes in cell width result in a transient increase in aspect ratio (*η* = *L/w*) during nutrient downshift, or a transient decrease in *η* during nutrient upshift (Fig. 2C). After nutrient shift, aspect ratio reaches its pre-stimulus homeostatic value over multiple generations. Typical timescale for transition to the new steady-state is controlled by the growth rate of the new medium (*∝ k^−^*^1^), such that the cell shape parameters reach a steady state faster in media with higher growth rate. This result is consistent with the experimental observation that newborn aspect ratio reaches equilibrium faster in fast growing media [6] (Figure 2—figure supplement 1F). In our model, cell shape changes are controlled by two parameters: the ratio *k/k*_*P*_ that determines cell aspect ratio, and *k/β* that controls cell width (Fig. 2D). Nutrient upshift or downshift only changes the ratio *k/β* while keeping the steady-state aspect ratio (*∝ k/k_P_*) constant.

We further used our model to predict drastic shape changes, leading to deviations from the homeostatic aspect ratio, when cells are perturbed by FtsZ knockdown, MreB depletion, and antibiotic treatments that induce non steady state filamentation (Fig. 2E). First, FtsZ depletion results in long cells while the width stays approximately constant [29]. We modelled FtsZ knockdown by decreasing *k*_*P*_ and simulations quantitatively agree with experimental data. Second, MreB depletion increases the cell width and slightly decreases cell length while keeping growth rate constant [29]. We modelled MreB knockdown by decreasing *β* as expected for disruption in cell wall synthesis machinery, while simultaneously increasing *k*_*P*_. This increase in *k*_*P*_ is consistent with a prior finding that in MreB mutant cells of various sizes, the total FtsZ scales with the cell width [31]. Third, transient long filamentous cells resulted from exposure to high dosages of cell-wall targeting antibiotics that prevent cell division, or DNA-targeting antibiotics that induce filamentation via SOS response [12]. Cell-wall targeting antibiotics inhibit the activity of essential septum forming penicillin binding proteins, preventing cell septation. We modelled this response as an effective reduction in *k*_*P*_, while slightly decreasing surface synthesis rate *β*. For DNA targeting antibiotics, FtsZ is directly sequestered during SOS response resulting in delayed ring formation and septation [33]. Surprisingly all filamentous cells have a similar aspect ratio of 11.0 *±* 1.4, represented by a single curve in the *S*-*V* plane (Fig. 2E).

The conserved surface-to-volume scaling in diverse bacterial species, *S ∼ V* ^2/3^, is a direct consequence of aspect-ratio homeostasis at the single-cell level. We present a regulatory model where aspect-ratio control is the consequence of a constant ratio between the rate of cell elongation (*k*) and division protein accumulation (*k*_*P*_). Deviation from the homeostatic aspect ratio is a consequence of altered *k/k*_*P*_, as observed in filamentous cells or MreB depleted cells. Aspect ratio control may have several adaptive benefits. For instance, increasing cell surface-to-volume ratio under low nutrient conditions can result in an increased nutrient influx to promote cell growth (Fig. 1C). Under translation inhibition by ribosome-targeting antibiotics, bacterial cells increase their volume while preserving aspect ratio [3, 9]. This leads to a reduction in surface-to-volume ratio to counter further antibiotic influx. Furthermore, recent studies have shown that the efficiency of swarming bacteria strongly depends on their aspect ratio [34, 35]. The highest foraging speed has been observed for aspect ratios in the range 4-6 [34], suggesting that the maintenance of an optimal aspect ratio may have evolutionary benefits for cell swarmers.

## Appendix

## Cell shape analysis

Bacterial cell surface area and volume are obtained directly from previous publications where these values were reported [3, 9, 14], or they are calculated assuming a sphero-cylindrical cell geometry using reported values for population-averaged cell length and width [12, 13, 17–20, 22, 23, 29]. For a spherocylinder of pole-to-pole length *L* and width *w*, the surface area is *S* = *wLπ*, and volume is given by 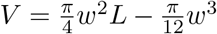. In the case of *S. aureus*, surface area and volume are computed assuming prolate spheroidal shape using reported population averaged values of cell major axis, *c*, and minor axis *a* [21]. Surface area of a prolate spheroid is 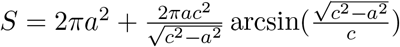, and volume is 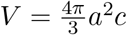.

## Initiator model for cell division

We considered a two-component model for FtsZ dynamics [28] - a cytoplasmic component with abundance *P*_c_, and a Z-ring bound component with abundance *P*_r_. Production of cytoplasmic and ring-bound FtsZ are given by: 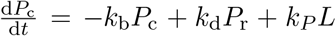, 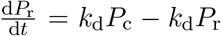, where *k*_*P*_ is the constant production rate of cytoplasmic FtsZ, *k*_b_ is the rate of binding of cytoplasmic FtsZ to the Z-ring, and *k*_d_ is the rate of disassembly of Z-ring bound FtsZ. At the start of the cell cycle, *P*_c_ = *P*^∗^ (a constant), *P*_*r*_ = 0, and the cell divides when the number of division proteins at the cell surface, *P*_r_, reaches the threshold amount *P*_0_ *∝ πw*. Net accumulation of division proteins, *P* = *P*_c_ + *P*_r_ *− P^∗^*, then follows the simple equation: 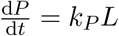, where *P* = 0 at the start of the cell cycle. Assuming *k*_b_ *» k*_d_ all the newly synthesized division proteins are recruited to the Z-ring such that cell division occurs when *P* = *P*_0_ (Fig. 2A). Upon division *P* is reset to 0. In the limit of FtsZ knockdown, septal ring formation occurs may be prevent, which can be realised in the limit that *k*_*b*_ is much smaller than the cell elongation rate.

## Cell growth simulations

We simulated the single-cell model using coupled equations for the dynamics of cell length (*L*), division protein (*P*) production, and cell width (*w*) (Fig. 2A) [18, 26]. In simulations, when *P* reaches the threshold, *P*_0_, the mother cell divides into two daughter cells whose lengths are 0.5 *± δ* fractions of the mother cell. Parameter *δ* is picked from Gaussian distribution (*µ* = 0, *σ* = 0.05). For nutrient shift simulations we simulated 10^5^ asynchronous cells growing with *k* = 0.75 h^*−*1^ (Fig. 2C). In Equation 3, parameter *β* = 4*k/w* is obtained from the fit to experimental data for 4*k/w* vs *k* (Figure 2—figure supplement 1B) [3]. At time 0 h we change *k* corresponding to nutrient upshift (*k* = 1.25, 2 h^*−*1^) or nutrient downshift (*k* = 0.75, 0.25 h^*−*1^). We calculated population average of length and width (Fig. 2C, middle), and population average of aspect ratio of newborn cells (Fig. 2C, right). Aspect ratio of newborn cells are binned in time and the bin average is calculated for a temporal bin size of 10 min. Examples of single cell traces during the nutrient shift are shown in Figure 2—figure supplement 1C-E. FtsZ depletion experiment [29] was simulated for *w* = 1 *µ*m while *k*_*P*_ was reduced to 40 % of its initial value. Best fit for MreB depletion experiment [29] was obtained for *η ≈* 2.7, by simulating reduction in division protein production rate, *k*_*P*_, and by varying *β* so that width spans range from 0.9 to 1.8 *µ*m. The best fit for long filamentous cells (resulting from DNA or cell-wall targeting antibiotics) was obtained for *η ≈* 11.0. Filamentation was simulated by decreasing *k*_*P*_ and *β* so that *w* spans the range from 0.9 to 1.4 *µ*m as experimentally observed [12].

## Acknowledgements

We thank Suckjoon Jun lab (UCSD) for providing single cell shape data for *E. coli*, and Javier López-Garrido, Guillaume Charras, and Deb Sankar Banerjee for useful comments. We gratefully acknowledge funding from EPSRC grant EP/R029822/1 (SB & NO), Royal Society Tata University Research Fellowship URF/R1/180187 (SB), Royal Society grant RGF/EA/181044 (SB & DS), and UCL Department of Physics & Astronomy (DS).

## Author Contributions

NO and SB conceived and designed research. NO and DS performed research. NO and SB wrote the paper.

---

**Figure 1 — figure supplement 1.**
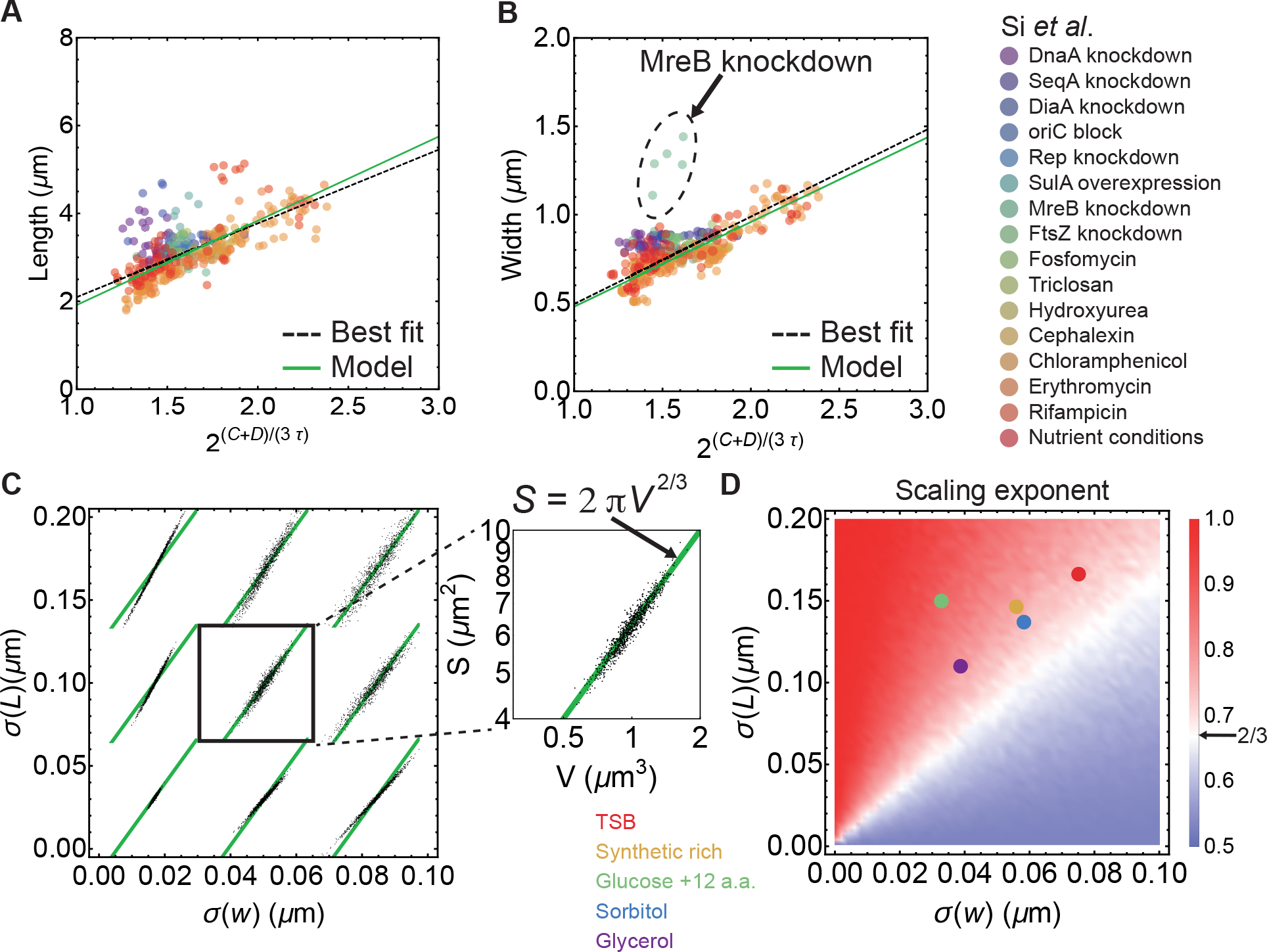
Control of cell width and length, and the influence of cell-to-cell variability. (A-B) Cell length and width vs 2^(*C*+*D*)*/τ*^ [3]. Here *C* is time from initiation to termination of DNA replication, *D* is time from termination of DNA replication to cell division, and *τ* is doubling time. Green lines are calculated assuming *S* = 2*πV* ^2/3^. Dashed black lines are best fit curves. (C) Variations in surface area and volume for a sphero-cylindrical bacterium, computed using normal distribution for cell widths (*µ*_*w*_,*σ*(*w*)) and log-normal distribution (*µ*_*L*_,*σ*(*L*)) of cell lengths. Green line is *S* = 2*πV* ^2/3^. (D) Surface-to-volume scaling exponent computed for different values of *σ*(*w*) and *σ*(*L*) while keeping *L/w* = 4. For each pair of values (*σ*(*w*), *σ*(*L*)) we pick 10^4^ random numbers from corresponding distributions and computed surface-to-volume scaling exponent. Total of 2500 pairs (*σ*(*w*), *σ*(*L*)) were used. We obtained *σ*(*w*) and *σ*(*L*) of newborn cells grown in mother machine by fitting experimental distributions. These values are shown by coloured points that correspond to different growth media (data from Taheri-Araghi *et al.* [6]). Cell-to-cell or intergenerational variability results in scaling exponents slightly above 2/3, as expected.

**Figure 2 — Figure supplement 1.**
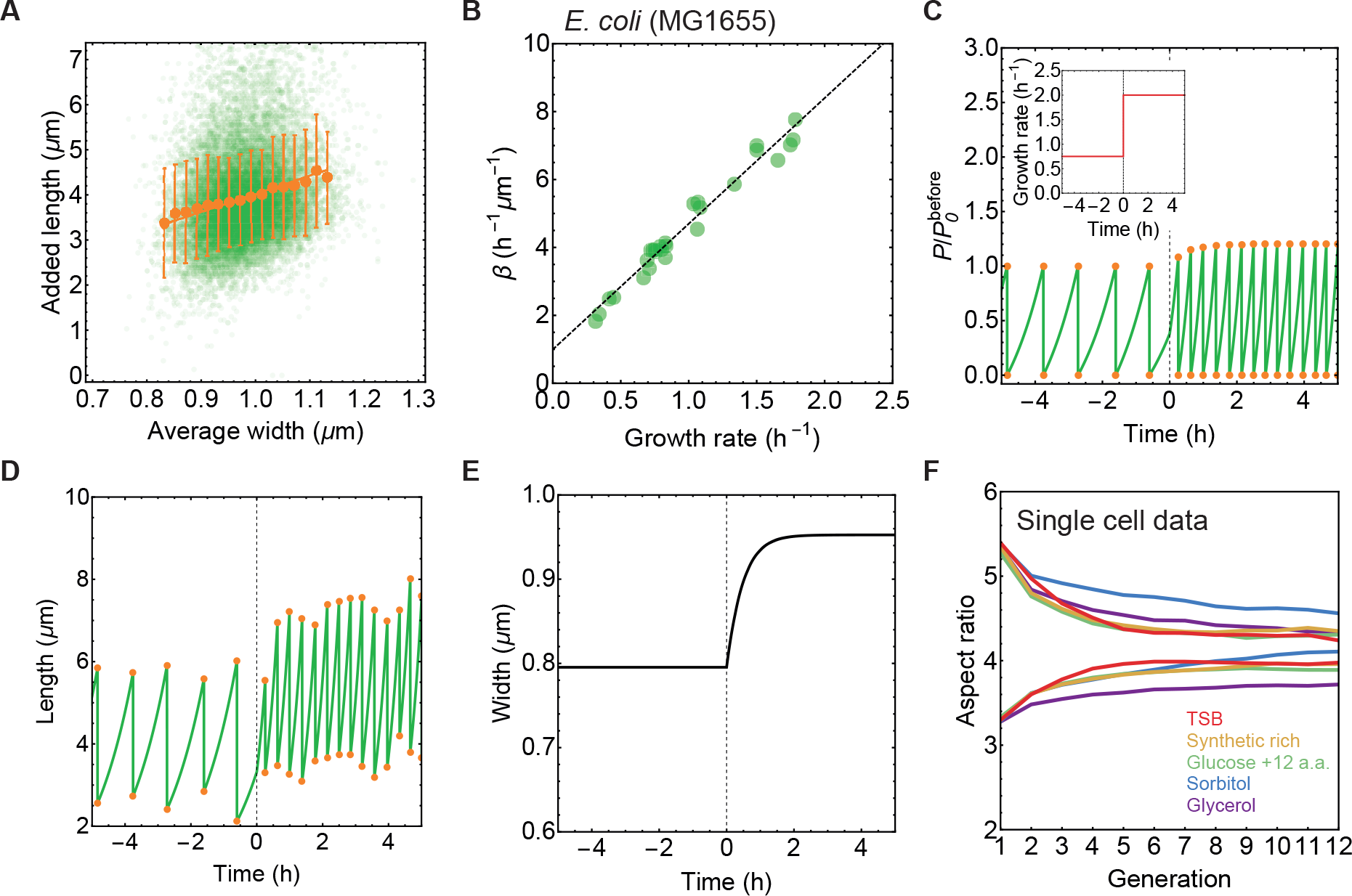
Single-cell data and simulations of nutrient upshift. (A) Single cell added length (∆*L*) vs average width (*w*) during one cell cycle for cell grown in TSB (Taheri-Araghi *et al.* [6]). Green circles represent single cell data, orange circles are average of binned data in width, error bars are *±*1 standard deviation, and orange line is best fit to binned data. (∆*L* = 4.014 *w*) (B) From experimental measurement of growth rate *k* and cell width *w* we calculated surface area production rate *β* = 4*k/w*. Data from Si *et al.* [3]. Dashed black line is best fit that we used in nutrient shift simulations. (C-E) Single cell traces for simulation of cell shape dynamics during nutrient upshift. (C) Division protein vs time normalised by *P*_0_ before the nutrient shift. (Inset) Growth rate vs time. (D) Length vs time. Length fluctuations at division is consequence of noise in division ratio (see Appendix). (E) Cell width vs time. (F) Aspect ratio of newborn cells vs generation number for single cell data from mother machine (Taheri-Araghi *et al.* [6]). Newborn cells with aspect ratio between 5-6 or 3-3.5 were tracked over generations. Population average for given generation number over 737-2843 cells for different growth condition is shown.

**Supplementary Table I.** Sample size for collected shape data [3, 6, 9, 12–14, 17, 19–23].

|  | Perturbations or Conditions | Number of cells per experiment or condition | Number of experiments or conditions |
| --- | --- | --- | --- |
| **Nonejuie et al.** | Antibiotics | ≥ 30 for elongated cell phenotype<br>≥ 50 for others | 44 |
| **Si et al.** | DnaA knockdown | 5205 - 9695 | 4 |
|  | SeqA knockdown | 4225 - 12034 | 8 |
|  | DiaA knockdown | ≥ 1000 | 8 |
|  | oriC block | 5939 - 10573 | 8 |
|  | Rep knockdown | 3144 - 5076 | 7 |
|  | SulA overexpression | 1215 - 6560 | 7 |
|  | MreB knockdown | 3718 - 18207 | 7 |
|  | FtsZ knockdown | ≥ 1000 | 8 |
|  | Fosfomycin | 1619 - 11170 | 7 |
|  | Triclosan | 4830 - 19082 | 7 |
|  | Hydroxyurea | 22158 - 36818 | 7 |
|  | Cephalexin | 2480 - 12429 | 8 |
|  | Chloramphenicol | 2202 - 21335 | 46 |
|  | Erythromycin | 5985 - 18955 | 16 |
|  | Rifampicin | 3458 - 10429 | 7 |
|  | Nutrient conditions | 2384 - 19566 | 15 |
| **Harris and Theriot** | Chloramphenicol | 82 | 5 |
|  | Fosfomycin | 190 | 5 |
| **Vadia et al.** | Nutrient conditions | 100 | 22 |
|  | Untreated | 100 | 3 |
|  | Rifampicin | 100 | 6 |
|  | Chloramphenicol | 100 | 5 |
|  | Cerulenin | 100 | 6 |
|  | Membrane synthesis | 100 | 21 |
| **Campos et al.** | Single deletion Keio | 174 - 408 | 4467 |
| **Taheri-Araghi et al.** | Glycerol | 14134 | 1 |
|  | Sorbitol | 10625 | 1 |
|  | Glucose + 12 a.a. | 18534 | 1 |
|  | Synthetic rich | 17574 | 1 |
|  | TSB | 18248 | 1 |
| **Harris and Theriot** | *C. crescentus* | 120 - 245 | 13 |
|  | *L. monocytogenes* | 113 - 185 | 7 |
| **Wright et al.** | *C. crescentus* | 250 | 1 |
| **Desmarais et al.** | *V. cholerae* | 1191 - 5448 | 5 |
|  | *P. aeruginosa* | 810 - 13065 | 6 |
| **Lopez-Garrido et al.** | *B. thuringiensis* | 20 - 30 | 1 |
|  | *B. pumilus* | 20 - 30 | 1 |
| **Ojkic et al.** | *B. subtilis* | 20 - 30 | 1 |
| **Carabetta et al.** | *B. subtilis* | 150 | 1 |
| **Quach et al.** | *S. aureus* | 12 - 71 | 406 |

